# The pseudobranch of jawed vertebrates is a mandibular arch-derived gill

**DOI:** 10.1101/2021.09.09.459574

**Authors:** Christine Hirschberger, J. Andrew Gillis

## Abstract

The pseudobranch is a gill-like epithelial elaboration that sits behind the jaw of most fishes. This structure was classically regarded as a vestige of the ancestral gill-arch like condition of the gnathostome jaw. However, more recently, hypotheses of jaw evolution by transformation of a gill arch have been challenged, and the pseudobranch has alternatively been considered a specialised derivative of the second (hyoid) pharyngeal arch. Here, we demonstrate by cell lineage tracing in a cartilaginous fish, the skate (*Leucoraja erinacea*), that the pseudobranch does, in fact, derive from the mandibular arch, and that it shares gene expression features and cell types with gills. We also show that the mandibular arch pseudobranch is supported by a spiracular cartilage that is patterned by a *shh*-expressing epithelial signalling centre. This closely parallels the condition seen in the gill arches, where cartilaginous appendages called branchial rays supporting the respiratory lamellae of the gills are patterned by a *shh*-expressing gill arch epithelial ridge (GAER). Taken together, these findings support serial homology of the pseudobranch and gills, and an ancestral origin of gill arch-like anatomical features from the gnathostome mandibular arch.

## Introduction

Hypotheses of jaw-gill arch serial homology are deeply rooted in the fields of vertebrate evolutionary biology and comparative anatomy and have profoundly shaped views on the evolutionary origin of the jawed vertebrate body plan. Apparent correspondence between the upper and lower jaw and the dorsal (epibranchial) and ventral (ceratobranchial) segments of the gill arch skeleton was recognised over a century ago and has led to various scenarios of jaw evolution by transformation of an anterior member of a hypothetical primitive series of gill-bearing arches (e.g., Gegenbaur, 1878; De Beer, 1937; Jarvik, 1980; Mallatt, 1996). There is now very strong evidence of a shared transcriptional network underlying axial patterning of the mandibular, hyoid and gill arch skeletons of jawed vertebrates (Gillis et al., 2013; Takechi et al., 2013; Hirschberger et al., 2021; reviewed by Medeiros and Crump, 2012), with some features of this network predating the origin of jaws (Cerny et al., 2010; Kuraku et al., 2010; reviewed by Square et al., 2017). These findings are consistent with serial homology of distinct dorsal, ventral, and intermediate domains of jawed vertebrate pharyngeal arches, and of the skeletal elements or tissues that derive, iteratively, from these domains.

The notion of an ancestral gill arch-like nature of the vertebrate mandibular arch, on the other hand, remains contentious. Extant jawless vertebrates (i.e., cyclostomes – lampreys and hagfishes) do not have a gill associated with their mandibular arch derivatives, and in a rare instance of preservation of putative gill impressions in the pharyngeal region of a stem-vertebrate fossil, such impressions are apparently lacking from the anterior-most arch (Conway Morris and Caron, 2014). Furthermore, the cyclostome mandibular arch gives rise to a velar skeleton that is morphologically distinct from the skeleton of the more caudal, gill-bearing arches (Johnels, 1948), and there is indirect evidence that some stem-gnathostomes may have possessed a similar pharyngeal endoskeletal organisation (Janvier, 1996; Janvier, 1981, 1985a,b as cited by Miyashita, 2016). Thus, in contrast to classical/neoclassical hypotheses of jaw origin by transformation of a gill arch, it has recently been suggested that the jaw instead evolved by “mandibular confinement” of an ancestrally distinct and gill-less oral apparatus (Miyashita, 2016). According to this hypothesis, parallels in gnathostome jaw and gill arch skeletal organisation reflect secondary assimilation of the former to the latter, rather than retention of a primitive similarity.

Unlike cyclostomes, however, many jawed vertebrates do possess a gill-like epithelial elaboration called a “pseudobranch” in close association with their mandibular arch derivatives. In teleost fishes, the pseudobranch resides in a subocular cavity (Laurent and Dunel-Erb, 1984), while in most elasmobranch cartilaginous fishes (sharks, skates, and rays) and some non-teleost actinopterygian bony fishes (e.g., sturgeons, paddlefishes, and bichirs), the pseudobranch resides within a spiracle – a small hole that opens behind the eye, and that permits sustained water intake into the buccopharyngeal cavity while the mouth is buried in the substrate or manipulating prey (Hughes, 1960) (Fig. 1A). During embryonic development, the spiracle forms by fusion of the first pharyngeal pouch with adjacent surface ectoderm and is therefore iteratively homologous with the gill slits that delineate the more caudal hyoid and gill arches. Furthermore, in elasmobranchs, the pseudobranch is supported by a spiracular cartilage, which sits in the anterior wall of the spiracle (Fig. 1B-C). This spiracular cartilage has been homologised with the cartilaginous branchial rays that project into interbranchial septa from the epi- and ceratobranchials of the gill arches, and which support the respiratory lamellae of the gills (El-Toubi, 1947; Gillis et al., 2009a).

**Figure 1:**
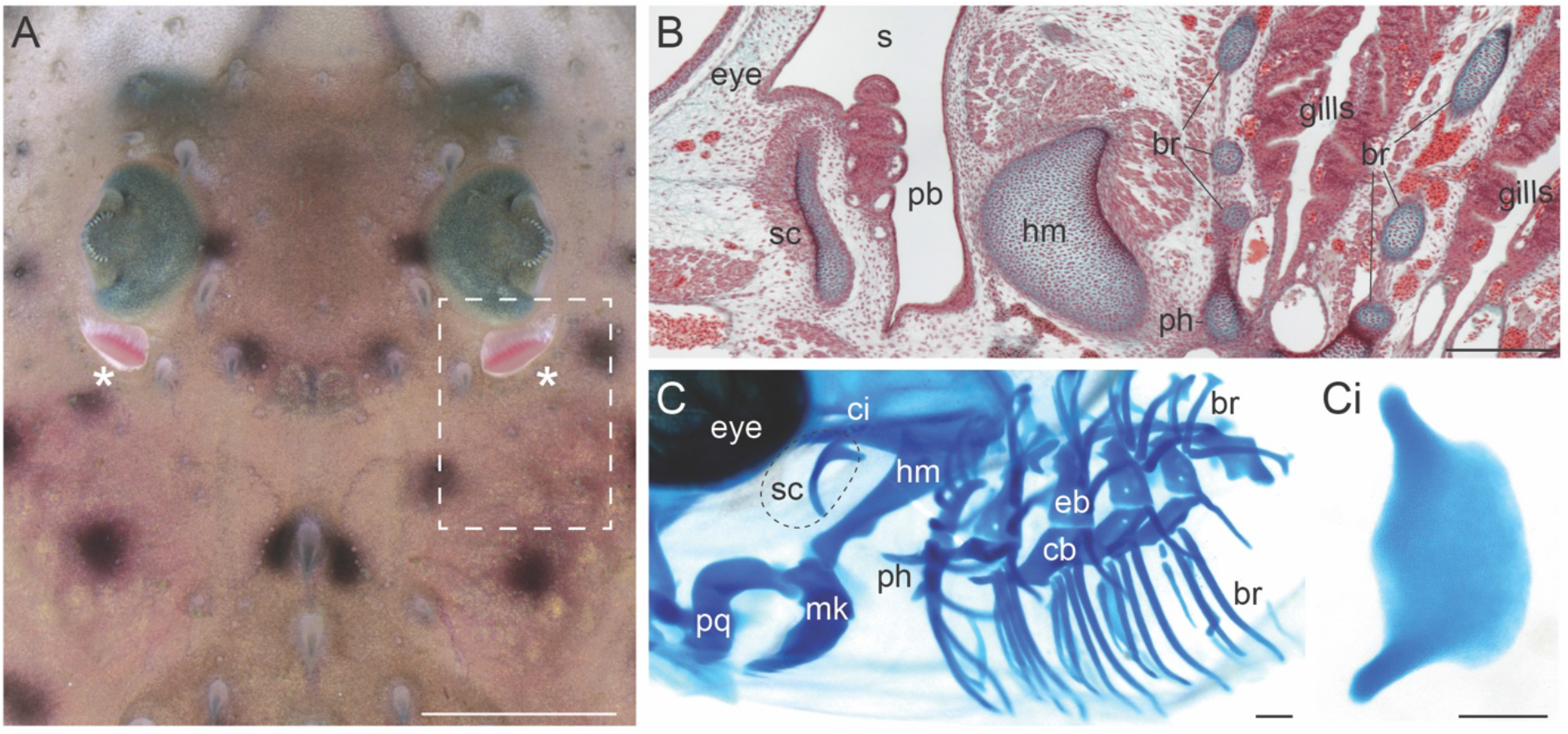
Anatomical overview of the spiracle, pseudobranch and spiracular cartilage in the little skate. **(A)** Dorsal view of the eye and spiracle of a skate hatchling. The spiracles are indicated with asterisks. Dashed box indicates region shown in **(B)** and **(C). (B)** Horizontal section of a S32 skate embryo showing the spiracular region. The pseudobranch sits inside the spiracle and is supported by the spiracular cartilage in the anterior spiracle wall. Stained with Masson’s trichrome. **(C)** Lateral view of a S32 skate embryo skeletal preparation showing the pharyngeal endoskeleton. The spiracular cartilage is located dorsal to the articular between the upper jaw (palatoquadrate) and the lower jaw (Meckel’s cartilage). **(Ci)** Frontal view of a dissected skate spiracular cartilage. The leaf-like sheet of cartilage supports the anterior wall of the spiracle. *br*, branchial rays; *eb*, epibranchials; *hm*, hyomandibula; *mk*, Meckel’s cartilage; *pb*, pseudobranch; *ph*, pseudohyal; *pq*, palatoquadrate; *s*, spiracle; *sc*, spiracular cartilage. Scale bars: A=3cm; B-Ci=500μm.

The close resemblance of the pseudobranch to the lamellae of the hyoid and gill arches, its location at the posterior margin of mandibular arch-derived structures and its formation, ancestrally, within a gill slit serial homologue have led to its interpretation as a reduced or vestigial mandibular arch-derived gill (Mallatt, 1996). But the pseudobranch also exhibits some features that could be regarded as atypical of a gill, mandibular arch-derived or otherwise. For instance, the pseudobranch is generally innervated by the facial (rather than trigeminal) nerve (Nilsson, 1984), and it receives oxygenated (rather than de-oxygenated) blood from the efferent hyoidean artery, via the afferent pseudobranchial artery (Hyrtl, 1838, as cited by Laurent and Dunel-Erb, 1984). For these reasons, it has been alternatively suggested that the pseudobranch is not a mandibular arch-derived gill, but rather is a specialised derivative of the second (hyoid) pharyngeal arch, which evolved to serve a non-respiratory, chemosensory function (Miyashita, 2016).

Here, we investigate the nature and development of the pseudobranch in a cartilaginous fish, the skate, *Leucoraja erinacea*. We demonstrate unequivocally, by histological and gene expression analyses and cell lineage tracing, that the skate pseudobranch is mandibular arch-derived, and that it shares cell types and gene expression patterns with the gill lamellae of the hyoid and gill arches. We further show that the spiracular cartilage, which supports the pseudobranch in skate, is patterned by a *shh*-expressing epithelial signalling centre, in a manner that closely parallels the way in which hyoid and gill arch branchial rays are patterned by a *shh*-expressing gill arch epithelia ridge (GAER). Taken together, our findings strongly support the pseudobranch as a mandibular arch-derived gill serial homologue and, we argue, as a vestige of an ancestral gill arch-like nature of the vertebrate mandibular arch.

## Methods

### Embryo collection and fate mapping

All animal work complied with protocols approved by the Institutional Animal Care and Use Committee at the Marine Biological Laboratory (MBL) in Woods Hole, U.S.A. Skate (*Leucoraja erinacea*) eggs were obtained from the Marine Resources Centre of the MBL. Staging was carried out according to Maxwell et al. (2008) and Ballard et al. (1993). Embryos for histological and gene expression analyses were fixed in 4% paraformaldehyde (PFA) in 1X phosphate-buffered saline (PBS) overnight at 4°C, the rinsed in 1X PBS, dehydrated into methanol and stored in methanol at -20°C prior to analysis. Embryos for cell lineage tracing experiments were maintained in a flow-through seawater system at ∼15°C. For CM-DiI labelling of the mandibular arch at S22, eggs were windowed, and microinjection of the mandibular arch with CM-DiI was performed *in ovo* according to Gillis et al. (2017). For labelling of the mandibular arch at S27, embryos were removed from the egg and anaesthetised in a petri dish of seawater containing 10mg/mL ethyl 3-aminobenzoate methanesulfonate salt (MS-222, Sigma) prior to microinjection according to Gillis and Hall (2016). CM-DiI-injected embryos were left to develop for 6-10 weeks, and then fixed in 4% PFA in PBS overnight at 4°C, rinsed three times in PBS and stored at 4°C in PBS with 0.01% sodium azide prior to analysis.

### Paraffin embedding and sectioning

All embryos for paraffin histology were dehydrated into 100% ethanol (or changed from 100% methanol to 100% ethanol), before clearing in Histosol (National Diagnostics). Embryos were washed 3 × 20 minutes in Histosol at room temperature, 2 × 30 minutes in 1:1 Histosol:paraffin at 60°C, and then then moved into molten paraffin for overnight at 60°C. The following day, embryos were washed 4 × 1 hour in paraffin, and then embedded in peel-a-way moulds. Embedded embryos were sectioned at 8μm on a Leica RM2125 rotary microtome, and sections were mounted on Superfrost Plus slides (Fisher).

### mRNA *in situ* hybridisation and immunofluorescence on paraffin sections

Chromogenic mRNA *in situ* hybridisation (ISH) for *foxl2* (MW457610), *gata2* (MZ501824), *shh* (EF100667) and *ptc* (EF100663) was performed in wholemount as in Hirschberger et al. (2021), and on paraffin sections as in O’Neill et al. (2007) with modifications according to Gillis et al. (2012). mRNA ISH by chain reaction (HCR) on paraffin sections was carried out according to Choi et al. (2018), with modifications as described in Criswell and Gillis (2020). Probe sets against *Leucoraja erinacea hif2a*/*epas1* (MZ501826; MI lot no. PRH337), *gata3* (MZ501825; MI lot no. PRJ403) and *gcm2* (MW389327; MI lot no. PRJ404) and hairpins were purchased from Molecular Instruments (https://www.molecularinstruments.com). Slides to be used for immunofluorescence were dewaxed in histosol and rehydrated through a descending ethanol series into 1X PBS + 0.1% Triton X-100 (PBT). For antigen retrieval, slides were preheated in distilled water for 5 minutes at 60°C, transferred to prewarmed antigen retrieval solution (10mM sodium citrate, pH6.0) and incubated for 25 minutes at 95°C. Slides were then cooled at -20°C for 30 minutes. Slides were rinsed 3 × 10 min in PBT, blocked for 30 min in 10% sheep serum and incubated in primary antibody under a parafilm coverslip in a humidified chamber overnight at 4°C. The next day, slides were rinsed 3 × 5 min in PBT and incubated in secondary antibody under a parafilm coverslip in a humidified chamber overnight at 4°C. Slides were then rinsed 3 × 10 min in PBT and coverslipped with Fluoromount G containing DAPI (Southern Biotech). Primary and secondary antibodies were diluted as follows: rabbit anti-5HT (Merck S5545, 1:250); mouse anti-SV2 (Developmental Studies Hybdridoma Bank, 1:100); goat anti-rabbit 488 (ThermoFisher A11008, 1:500); goat anti-mouse 633 (ThermoFisher A21050, 1:500).

### *in ovo* cyclopamine treatment, wholemount skeletal preparation and statistical analysis

*in ovo* cyclopamine treatment was performed according to Gillis and Hall (2016). Briefly, skate eggs have an approximate internal chamber volume of 12ml. To achieve an *in ovo* cyclopamine concentration of ∼20μM, 25μl of 9mM stock solution of cyclopamine in dimethylsulfoxide (DMSO) was injected into skate eggs at S22, S27 or S29, using a syringe and 30-gauge needle. For control embryos, an equivalent volume of DMSO was injected into the egg. Cyclopamine-treated embryos were grown to S32, and surviving embryos (n=6/21, n=12/28, n=7/19 for cyclopamine treatment at S22, S27 and S29 respectively, and n=3/3, n=4/5 and n=5/5 for DMSO control treatment at S22, S27 and S29 respectively) were fixed as described above and analysed for skeletal defects by skeletal preparation.

For wholemount staining of the endoskeleton, skate embryos were moved from 100% methanol into 70% ethanol at room temperature. Embryos were then stained with 0.2mg/mL Alcian Blue 8GX in acetic ethanol (3:7 glacial acetic acid: absolute ethanol) for 24 hours with gentle rocking at room temperature. Embryos were destained in acetic ethanol for 24 hours, rehydrated through a descending ethanol series, washed in ddHOH, and digested in 1% w/v trypsin in 2% w/v sodium tetraborate for 20 minutes. Embryos were then cleared in a graded 0.5% w/v KOH:glycerol series (3:1, 1:1, 1:3 0.5% KOH:glycerol) for 24hrs/step. Specimens were stored in 80% glycerol at 4°C and manually dissected for photography.

For the statistical analysis, the dissected left spiracular cartilage of each embryo was imaged on a Leica M165FC stereomicroscope, and the surface area of each cartilage was measured using Fiji (Schindelin et al., 2012) and normalised against embryo total length. To test for statistically significant differences among the means of normalised spiracular cartilage area for each group (DMSO at S22, cyclopamine at S22, DMSO at S27, cyclopamine at S27, DMSO at S29 and cyclopamine at S29), a Kruskal-Wallis analysis of variance was performed, followed by a Dunn’s test with Bonferroni correction to identify which pairwise comparisons were statistically significantly different. The same statistical analysis confirmed no significant difference between normalised spiracular cartilage area of DMSO-treated vs. wild-type (n=3) spiracular cartilages at S32 (p=0.305).

## Results

### The skate pseudobranch is mandibular arch-derived

In the skate, the developing pseudobranch is first apparent at S29, as a thickening of the epithelium of the posterior mandibular arch (Fig. 2A), and by S30 we observe the first evidence of vasculature subjacent to this thickening (Fig. 2B). By early S31 (Fig. 2C), the pseudobranch is a gill lamella-like structure that extends into the spiracular cavity from the posterior mandibular arch, and within the mandibular arch, we see condensation of the spiracular cartilage. The pseudobranch and spiracular cartilage continue to develop through later S31 (Fig. 2D), and by S32, these structures appear fully differentiated (Fig. 2E). Although the spiracular cartilage is closely associate the palatoquadrate (Fig. 1), these elements condense and chondrify entirely independently of one another (Fig. 2F-H).

**Figure 2:**
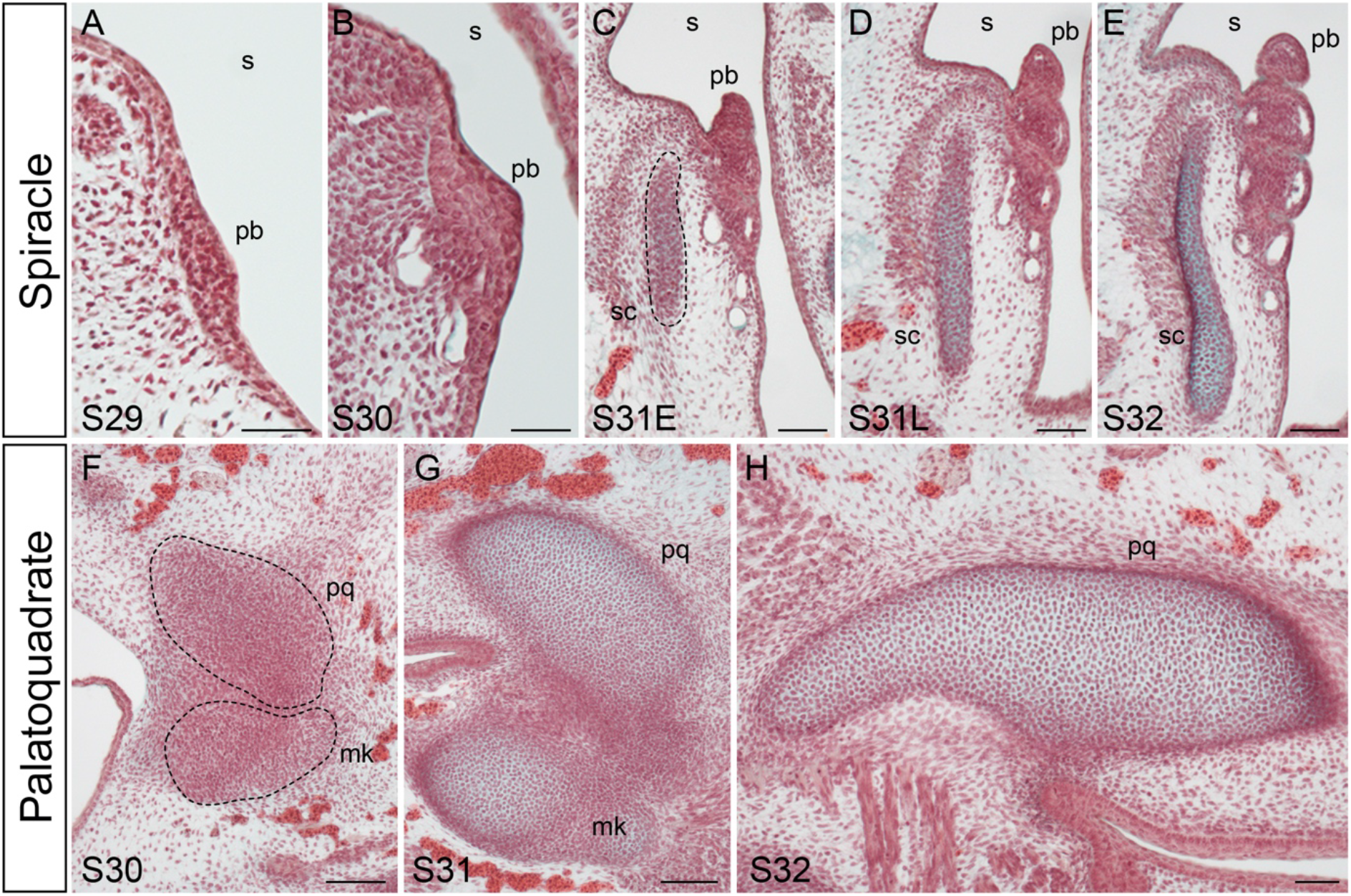
Development of the pseudobranch, spiracular cartilage and palatoquadrate in the little skate. **(A)** At S29, the development of the pseudobranch is first apparent as a thickening of the posterior epithelium of the mandibular arch. **(B)** By S30, the pseudobranch thickens, further, and vasculature lies adjacent to it. **(C-D)** At S31, the pseudobranch is extending from the posterior epithelium of the mandibular arch and has taken on a lamellar-like morphology, and the spiracular cartilage is condensing within subjacent mesenchyme (dashed line in **C**). **(E)** By S32, the pseudobranch and spiracular cartilage have fully differentiated. Note that the spiracular cartilage chondrifies completely independently of the **(F)** condensing and **(G-H)** differentiating palatoquadrate. *mk*, Meckel’s cartilage; *pb*, pseudobranch; *pq*, palatoquadrate; *s*, spiracle; *sc*, spiracular cartilage. Scale bars: A-H=50μm.

Although the pseudobranch clearly appears to develop from the posterior side of the mandibular arch in the skate, we sought to directly confirm this embryonic origin by cell lineage tracing. We and others have previously reported that *foxl2* is expressed in the core mesoderm and presumptive gill-forming epithelium of the pharyngeal arches of cartilaginous fishes (Wotton et al., 2007; Hirschberger et al., 2021). Analysis of *foxl2* expression by mRNA *in situ* hybridisation (ISH) in skate embryos at S24 reveals these expression patterns not only in the developing hyoid and gill arches, but also in the developing mandibular arch (Fig. 3A, B). We speculated that posterior mandibular arch epithelial expression of *foxl2* was serially homologous with expression in the developing gill epithelium of the hyoid and gill arches and marked the presumptive pseudobranch epithelium.

**Figure 3:**
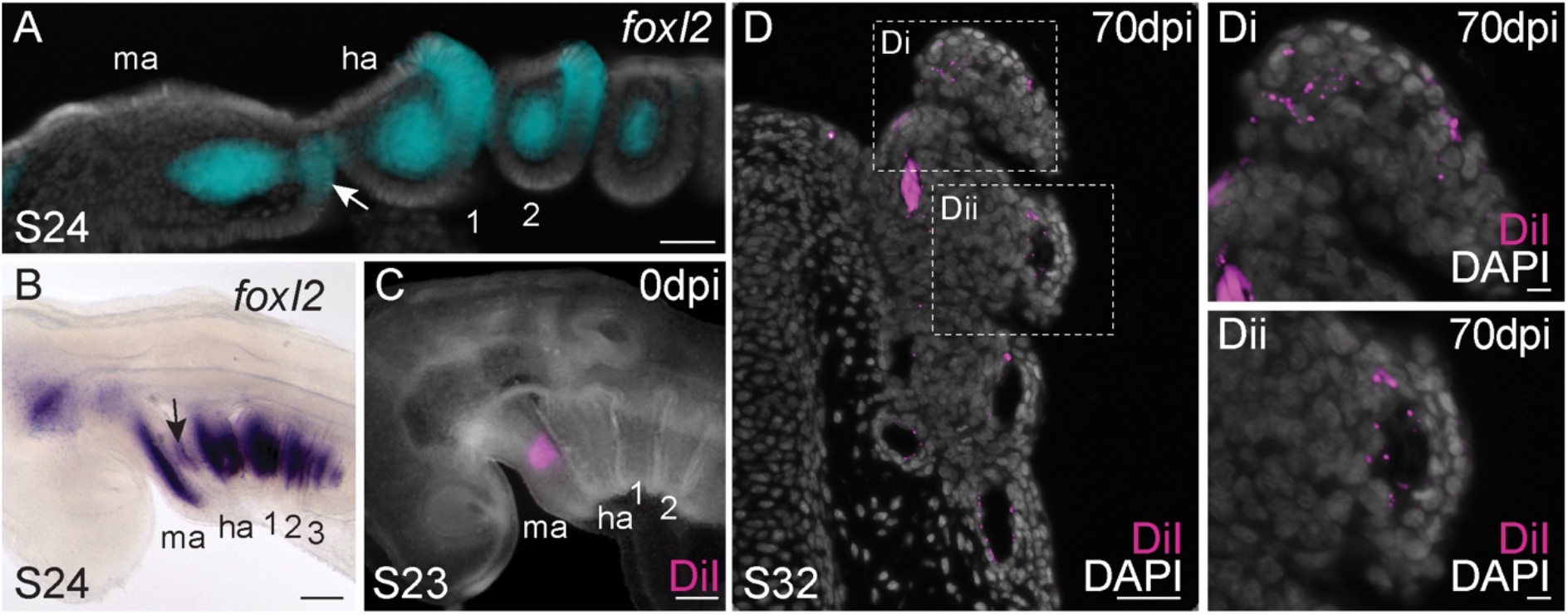
The skate pseudobranch is mandibular-arch-derived. **(A)** At S24, *foxl2* is expressed in the mesodermal core of all pharyngeal arches, as well as in the presumptive pseudobranch and gill endodermal epithelium of the mandibular (white arrow) and hyoid/gill arches, respectively. **(B)** *foxl2*+ presumptive pseudobranch epithelium (black arrow) is also apparent on the posterior mandibular arch following wholemount mRNA ISH. **(C)** Microinjection of the lipophilic dye CM-DiI in the posterior dorsal-intermediate mandibular arch of skate embryos at S23 to broadly label mandibular arch derivatives. **(D)** By S32 (after ∼10 weeks of post-injection development), CM-DiI-positive cells are recovered throughout the skate pseudobranch, including **(Di)** within the lamellae of the pseudobranch as well as **(Dii)** in the walls of the pseudobranch vasculature. *ha*, hyoid arch; *ma*, mandibular arch; *sc*, spiracular cartilage; *1-3*: gill arches 1-3. Scale bars: A-D=50μm; Di, Dii = 5μm.

To test this, we broadly labelled the dorsal-intermediate mandibular arch in skate embryos at S23 with the lipophilic dye CM-DiI (Fig. 3C). At S23, the first pharyngeal endodermal pouch has fully fused with the surface ectoderm, permitting us to label mandibular arch tissues broadly and specifically without risk of contaminating the hyoid arch. Injected embryos were then reared until S32, by which point the pseudobranch has fully differentiated. Of 11 surviving embryos, 9 retained CM-DiI-positive cells throughout the entirety of the pseudobranch (Fig. 3D). CM-DiI-positive cells were recovered throughout the pseudobranch lamellae (Fig. 3Di), as well as in the endothelial lining of the pseudobranch vasculature (Fig. 3Dii). No CM-DiI positive cells were recovered in any hyoid arch-derived features. These gene expression and lineage tracing findings conclusively demonstrate the mandibular arch origin of the pseudobranch in skate.

### Conservation of gene expression between the skate pseudobranch and gill lamellae

The gill lamellae of bony fishes and cyclostomes possess neuroepithelial cells (NECs), which function to mediate physiological responses to hypoxia, and which are identifiable by their immunoreactivity for serotonin (5-HT) and synaptic vesicle glycoprotein 2 (SV2) (Dunel-Erb et al., 1982; Jonz and Nurse, 2003, 2005; Hockman et al., 2017). In skate gills at S32 (Fig. 4A), immunofluorescent detection of 5HT and SV2 confirms conservation of this putative hypoxia-sensitive neuroepithelial cell type in the tips of the gill lamellae (Fig. 4B, 4Bi-iii). Using mRNA ISH, we also find that differentiated skate gill lamellae express the transcription factors *epas1*/*hif2a, gata2, gata3, gcm2 and foxl2* (Fig. 4C-E; Fig. S1A-B), with *epas1*/*hif2a* expression likely reflecting cells that function in hypoxia-sensitivity and/or stress response (Semenza, 2000; Dioum et al., 2009), and *gata2/3* and *gcm2* expression reflecting putative homology between the gills of fishes and the amniote parathyroid gland (Okabe and Graham, 2004).

**Figure 4:**
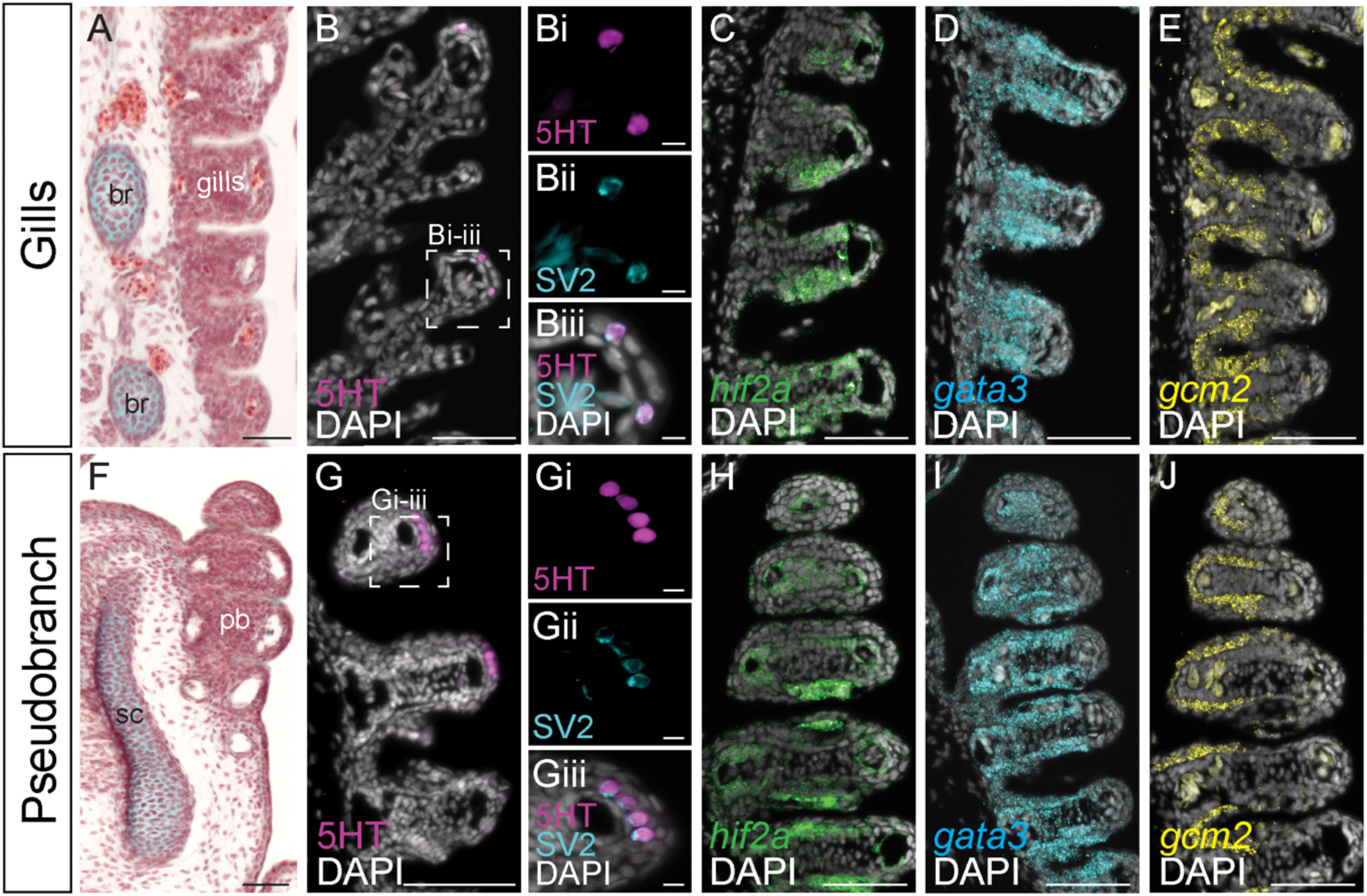
Shared gene expression features of gills and the pseudobranch in skate. **(A)** In skate, interbranchial septa bearing gill lamellae are internally supported by cartilaginous branchial rays. **(B)** Putative hypoxia-sensitive neuroepithelial cells (NECs) co-expressing **(Bi-Biii)** serotonin and SV2 are present in S32 skate gill lamellae. **(C)** Skate gills are marked by the expression of *hif2a*, **(D)** *gata3* and **(E)** *gcm2*. **(F)** The skate pseudobranch is internally supported by the spiracular cartilage. **(G)** NECs co-expressing **(Gi-Giii)** serotonin and SV2 are also present in the lamellae of the skate pseudobranch. **(H)** The skate pseudobranch shares expression of *hif2a*, **(I)** *gata3* and **(J)** *gcm2* with gills. *br*, branchial rays; *sc*, spiracular cartilage; *pb*, pseudobranch. Scale bars: A,B, C-D, F, G, H-I=50μm, Bi-Biii, Gi-Giii=5μm.

Immunostaining and ISH on sections revealed conservation of the above gene expression features in the skate pseudobranch (Fig. 4F). We found 5HT/SV2+ NECs in the tips of the pseudobranch lamellae (Fig. 4G, Gi-Giii), as well as pseudobranch expression of *hif2a, gata2, gata3, gcm2* and *foxl2* (Fig. 4H-J; Fig. S1C-D). These observations point to shared cell types and gene expression features – and, by extension, function – between the differentiated pseudobranch and gills of the skate, consistent with the serial homology of these structures.

### Common origin of the skate spiracular cartilage and branchial rays from serially homologous domains of sonic hedgehog-responsive mesenchyme

The cartilaginous branchial rays of that support the gill lamellae of the hyoid and gill arches of skates and sharks are patterned by a *shh*-expressing epithelial signalling centre called the gill arch epithelial ridge (GAER) (Gillis et al., 2009b, Gillis et al., 2011; Gillis & Hall, 2016). In these arches, *shh* is initially broadly expressed in the posterior arch epithelium, and this expression then resolves into the GAER as the arches undergo a lateral expansion. Cell lineage tracing has shown that branchial rays develop from GAER *shh*-responsive mesenchyme, and inhibition of GAER shh signalling results in a truncation/reduction of the branchial ray skeleton (Gillis et al., 2009b; Gillis & Hall, 2016).

We characterised *shh* expression in the developing mandibular arch of the skate. We found that at S22, *shh* was broadly expressed in the posterior epithelium of the skate mandibular arch (Fig. 5A-B), with this expression resolving into a GAER-like ridge of *shh*-expressing cells along the posterior margin of the mandibular arch by S27 (Fig. 5C-D). This ridge of *shh* expression persisted into S29, when it delineated the anterior edge of the spiracle (Fig. 5E-F). These patterns of *shh* expression in the mandibular arch ae indistinguishable from the GAER *shh* expression of the branchial ray-bearing hyoid and gill arches and led us to speculate about their possible skeletal patterning functions within the mandibular arch.

**Figure 5:**
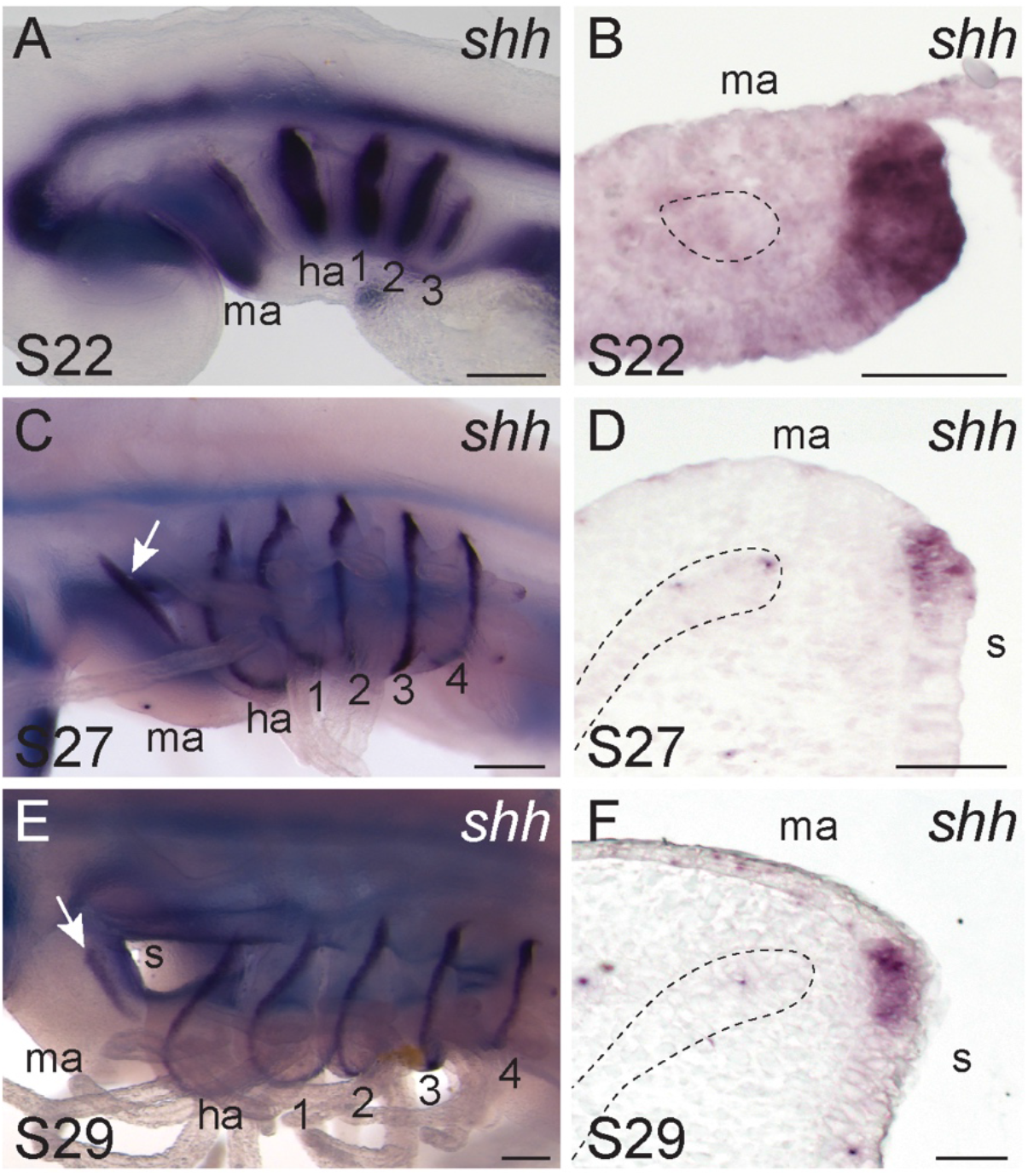
A *shh*-expressing epithelial ridge on the skate mandibular arch. **(A)** At S22, *shh* is expressed broadly throughout the posterior epithelium of each pharyngeal arch in skate, including **(B)** the mandibular arch. **(C)** By S27, *shh* expression on the skate mandibular arch has resolved to a **(D)** thin ridge of cells lying along the posterior-lateral edge of the arch, comparable to the gill arch epithelial ridge (GAER) *shh* expression of the hyoid and gill arches. **(E**,**F)** At S29, *shh* persists in the mandibular hyoid and gill arch GAERs. In **(B), (D)** and **(F)**, the mesodermal core or muscle plate of the mandibular arch is outlined with a dashed line. *ha*, hyoid arch; *ma*, mandibular arch; *1-4*, gill arches 1-4. Scale bars: A, C, E=200μm, B, D, F=50μm.

To test the fate of mesenchyme underneath the *shh*-expressing GAER of the skate mandibular arch, we labelled this tissue by microinjection of CM-DiI in skate embryos at S27 (Fig. 6A). We laeled *ptc*+ (shh-responsive) mesenchymal cells underneath the mandibular arch GAER (Fig. 6B-D), reared injected embryos until S32, and then assessed the contribution of that tissue to the mandibular arch skeleton. Histological analysis of labelled embryos revealed CM-DiI-positive cells throughout the spiracular cartilage (Fig. 6E,F; Fig. S2; n=22/29), as well as in the connective tissue surrounding the spiracular cartilage. These experiments indicate that, like gill arch branchial rays, the spiracular cartilage develops from GAER shh-responsive mesenchyme of the mandibular arch.

**Figure 6:**
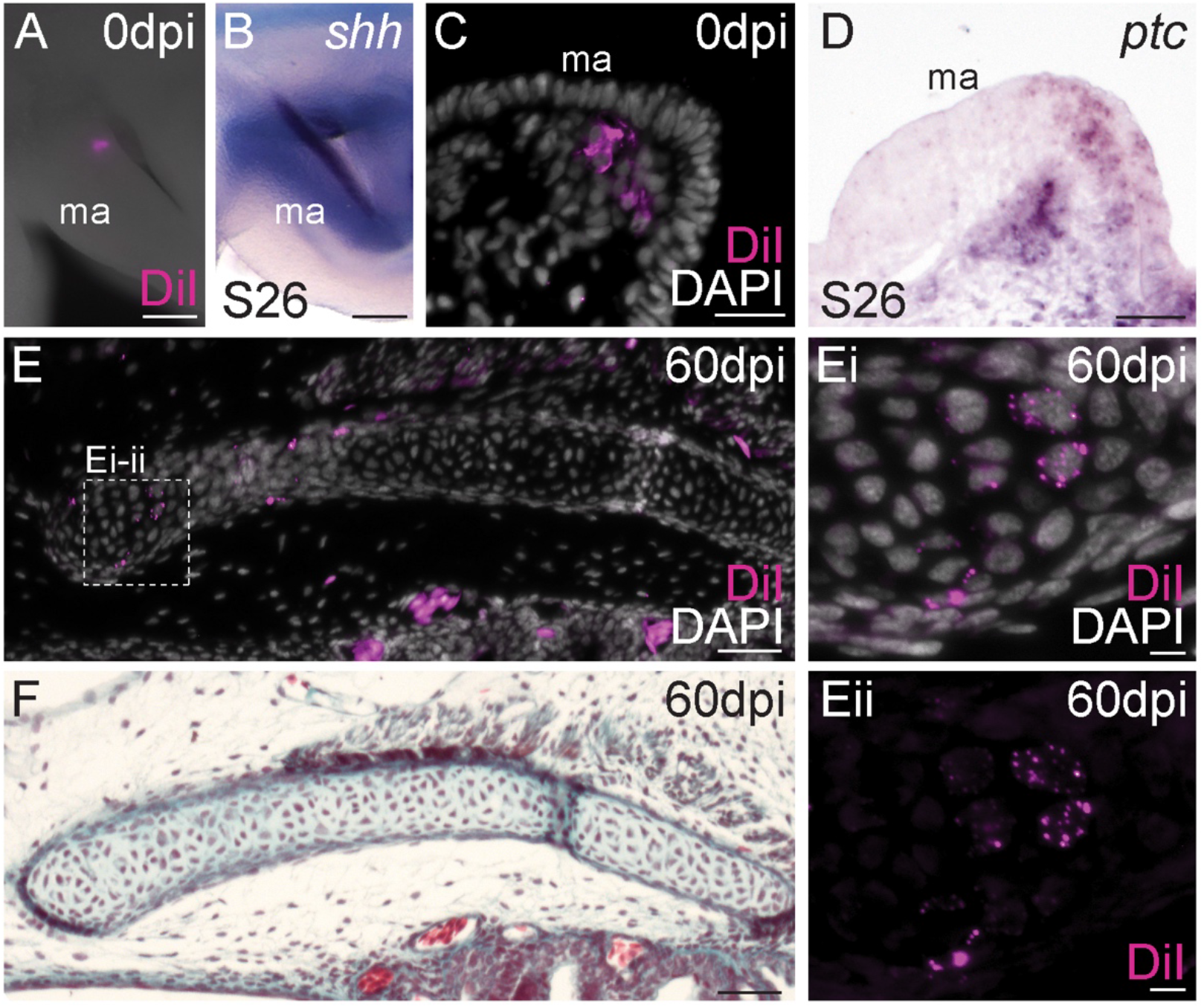
The skate spiracular cartilage derives from GAER shh-responsive mandibular arch mesenchyme. **(A)** Microinjection of the lipophilic dye CM-DiI in the mandibular arch mesenchyme **(B)** subjacent to the *shh*-expressing GAER results in **(C**,**D)** focal labelling of *ptc*+ (i.e. shh-responsive) mesenchyme of the posterior mandibular arch. **(E)** At 60 days post-injection, CM-DiI-positive chondrocytes are recovered throughout the **(F)** spiracular cartilage. *ma*, mandibular arch. Scale bars: A-F=50μm, Ei, Eii=5μm.

### The skate spiracular cartilage and branchial rays share shh dependent patterning mechanisms

Finally, we set out to test whether mandibular arch GAER shh signalling patterns the spiracular cartilage in a manner that is comparable to the GAER patterning of hyoid and gill arch branchial rays. We have previously shown in skate that from S22-27, successively earlier inhibition of hedgehog signalling by cyclopamine treatment resulted in a progressively greater reduction of gill arch mesenchymal cell proliferation, and a corresponding reduction in the number of branchial rays on each arch (Gillis & Hall, 2016). To test the effects of hedgehog signalling inhibition on development of the spiracular cartilage, we conducted *in ovo* cyclopamine treatments with skate embryos at S22, S27 and S29, as per Gillis and Hall (2016), and assessed surviving embryos for spiracular cartilage defects at S32. Given the 2-dimensional sheet-like nature of the skate spiracular cartilage, our test for spiracular cartilage reduction consisted of skeletal preparation, unilateral (left) spiracular cartilage dissection, measurement of spiracular cartilage area (standardised against total embryo length), and pairwise comparisons of mean ratios between control and cyclopamine-treated embryo at each stage.

Control (DMSO-treated) skate embryos showed no observable spiracular cartilage defects (Fig. 7A, A’), and spiracular cartilage area was not significantly different from wildtype S32 embryos (n = 3, not shown). Conversely, cyclopamine treatment at S22 and S27 led to striking spiracular cartilage defects (Fig. 7B, B’, C, C’), and to an overall significant reduction in spiracular cartilage area, relative to control (Fig. 7E). After treatment at S22, spiracular cartilages were either completely absent (n=3/6) or present as substantially reduced nodules of cartilage (n=3/6), while defects in embryos treated at S27 ranged from complete loss of the spiracular cartilage (n=2/12) to a reduction of the characteristic leaf-like sheet of cartilage, leaving only the slender endpoints connected by a thin bridge of cartilage (n=10/12). In contrast, cyclopamine treatment at S29 resulted in spiracular cartilages with some slightly abnormal morphology (Fig. 7D, D’), but no significant reduction in overall spiracular cartilage area relative to control (Fig. 7E). These findings reveal a shared role for GAER shh signalling in patterning the spiracular cartilage and branchial rays, and a shared period in development during which shh signalling is required for development of the spiracular cartilage and branchial rays.

**Figure 7:**
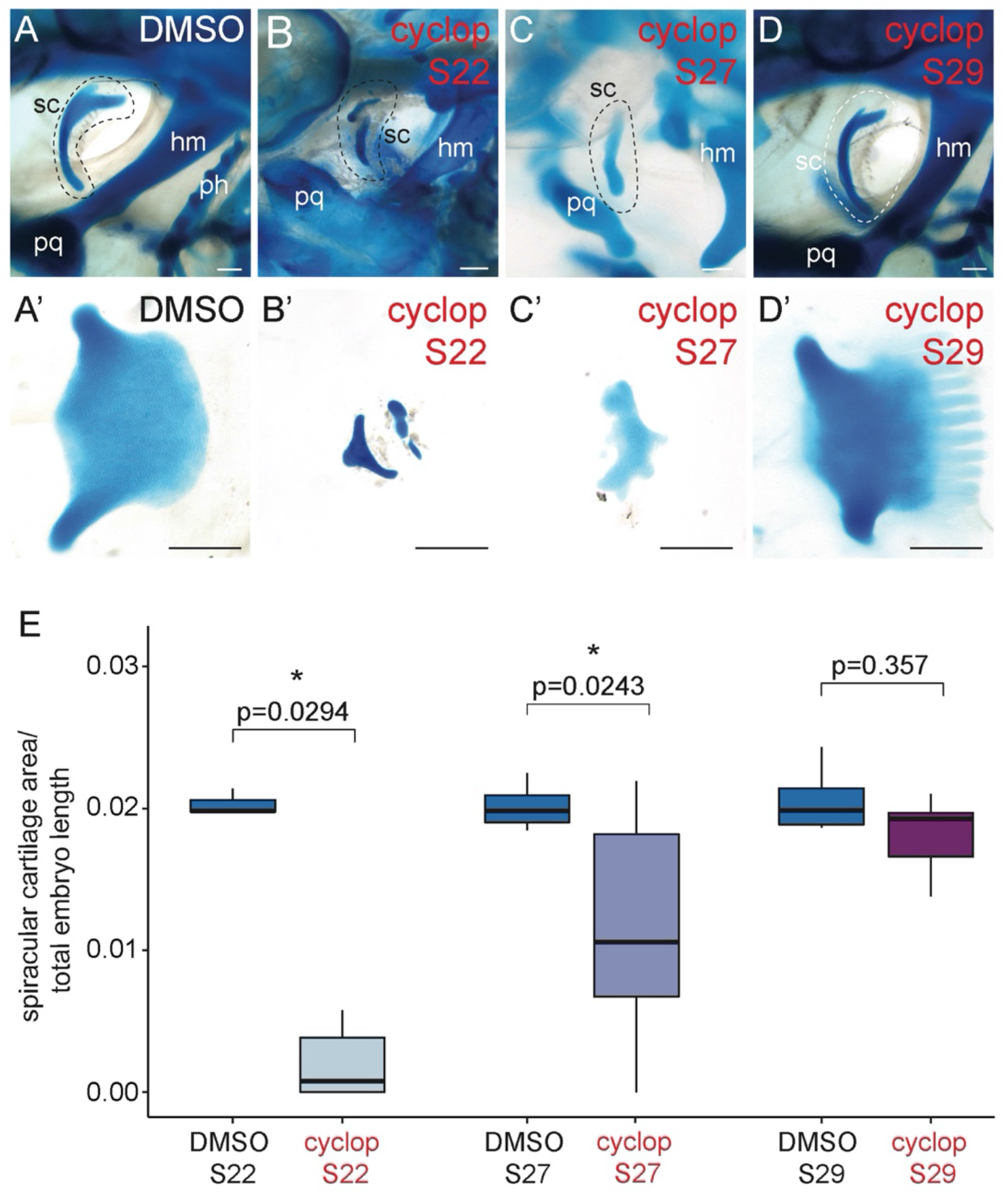
Patterning of the skate spiracular cartilage by shh signalling. **(A-A’)** DMSO (control) treatment *in ovo* has no effect on spiracular cartilage morphology, while treatment with 20uM cyclopamine *in ovo* at **(B-B’)** S22 and **(C-C’)** S27 causes spiracular cartilage malformation and reduction in spiracular cartilage size. **(D-D’)** Treatment with 20uM cyclopamine *in ovo* at S29 has no significant effect on spiracular cartilage size. **(E)** Analysis of spiracular cartilage size (spiracular cartilage area/total embryo length) between cyclopamine- and DMSO-treated skate embryos at S22, S27 and S29 reveals statistically significant reductions in spiracular cartilage size with cyclopamine treatment at S22 and S27 are significant, but no significant reduction in spiracular cartilage size with cyclopamine treatment at S29. Kruskal-Wallis test followed by Dunn’s test with Bonferroni correction. *hm*, hyomandibula; *ph*, pseudohyal; *pq*, palatoquadrate; *sc*, spiracular cartilage. Scale bars = 500μm.

## Discussion

Anatomical and histological similarities between the pseudobranch and gills are striking and have led to hypotheses of their homology – and, by extension, of an ancestrally gill-bearing mandibular arch (Romer, 1970; Mallatt, 1996). The latter remains contentious, owing to an absence of gill-like structures associated with mandibular arch derivatives in extant jawless vertebrates, and a lack of direct fossil evidence of the nature of first (mandibular) pharyngeal arch derivatives along the vertebrate and early gnathostome stems (Janvier, 1996; Miyashita, 2016). We have now presented unequivocal evidence for the mandibular arch origin of the pseudobranch in the skate, as well developmental, gene expression and patterning similarities shared by the pseudobranch and gills (and, in the skate, their supporting skeletal elements). We argue that these similarities may only reasonably be interpreted as products of serial homology – in this instance, not merely anatomical similarity due to “co-option” of a conserved patterning programme, but rather due to the derivation of the pseudobranch and gills from a primitive series of gill-bearing arches – and that the pseudobranch should, therefore, be interpreted as a vestigial mandibular arch-derived gill (Fig. 8).

**Figure 8:**
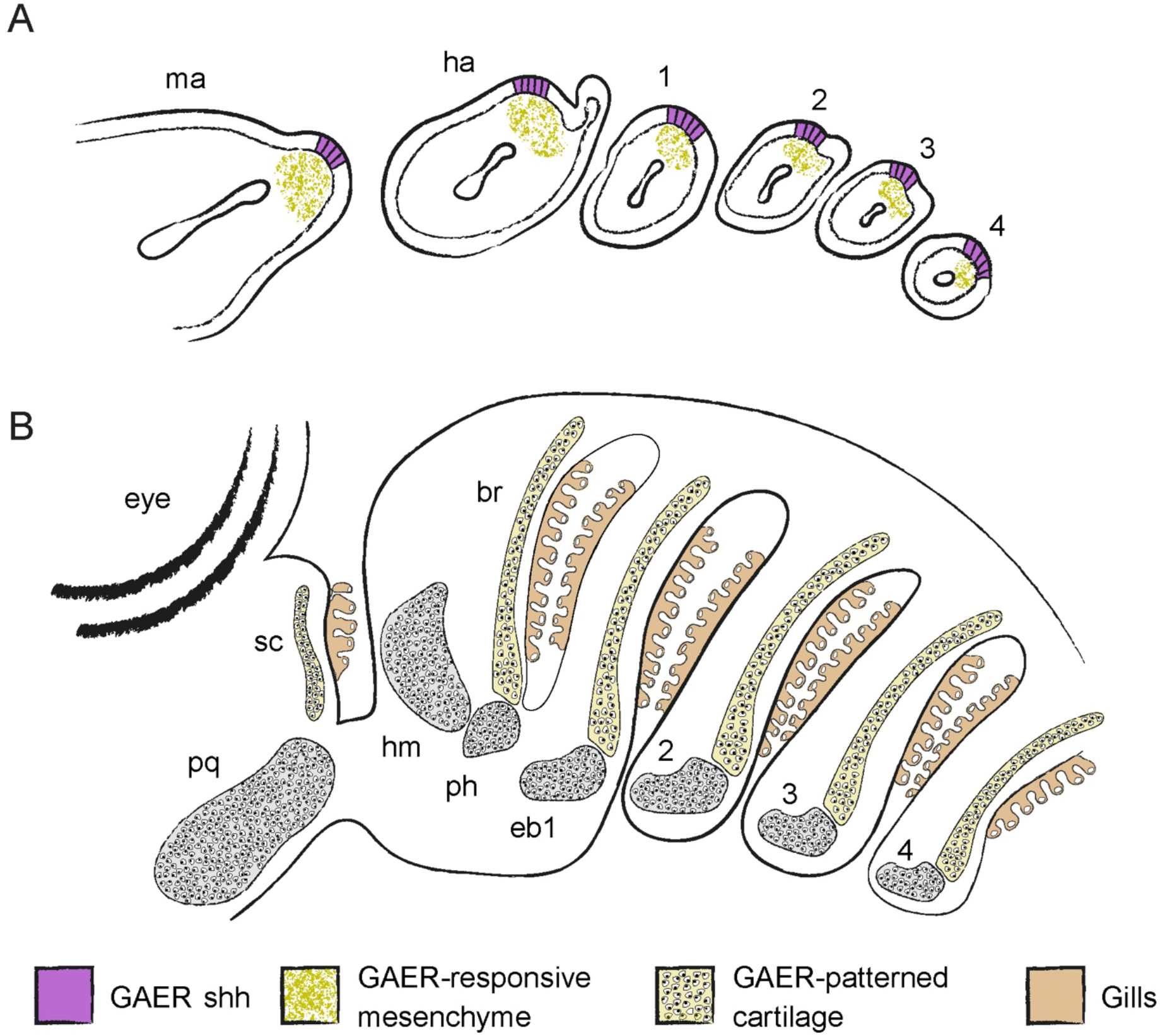
Serial homology of the mandibular arch pseudobranch and spiracular cartilage with the gill lamellae and branchial rays in the little skate. The mandibular, hyoid and gill arches of skate embryos each possess a *shh-*expressing gill arch epithelial ridge (GAER), which signals to underlying skeletogenic mesenchyme. These serially homologous signalling centres pattern the spiracular cartilage and branchial rays of the mandibular and hyoid/gill arches, respectively. The spiracular cartilage and branchial rays serve to support the serially homologous pseudobranch (mandibular arch) and gill lamellae (hyoid and gill arches), respectively. *br*, branchial rays; *eb1-4*, epibranchials 1-4; *ha*, hyoid arch; *hm*; hyomandibula; *ma*, mandibular arch; *ph*, pseudohyal; *pq*, palatoquadrate; *sc*, spiracular cartilage.

### Vasculature, innervation and embryonic origin of the pseudobranch

It has been argued, based on irrigation with oxygenated blood from the afferent hyoidean artery and innervation by the facial nerve, that the pseudobranch of gnathostomes is not a derivative of the mandibular arch, but rather of the hyoid arch (Miyashita, 2016). We have shown that, in fact, the pseudobranch derives from a domain of *foxl2*-expressing epithelium on the posterior margin of the mandibular arch in skate (just as hyoid and gill lamellae derive from *foxl2*-expressing epithelial domains of their respective pharyngeal arches). However, regardless of this direct evidence of embryonic origin, the vascular and neural characteristics described above needn’t necessarily preclude a mandibular arch origin of the pseudobranch. In most fishes, blood supply to the efferent pseudobranchial artery does derive from the afferent hyoidean in the adult condition, but this configuration arises from an embryonic one in which the vasculature of the pseudobranch more closely resembles that of the gills (Laurent and Dunel-Erb, 1984). For example, in shark embryos, the pseudobranch initially receives venous blood from the ventral aorta, with this connection subsequently degenerating as a novel blood supply from the afferent hyoidean artery is secondarily formed (De Beer, 1924; Daniel, 1934; Laurent and Dunel-Erb, 1984). Also, while mandibular arch-derived structures do tend to be innervated by the trigeminal nerve, there are exceptions: for example, the maxillary barbels of teleosts develop from the mandibular arch but are innervated by the facial nerve (Kanwal et al., 1987; Caprio et al., 2015; LeClair and Topczewski, 2010; Moore et al., 2012). These observations underscore the difficulty of ascertaining embryonic origin based on adult innervation or vasculature, and the importance of direct embryonic observation and cell lineage tracing for this purpose.

### Conservation of pseudobranch and gill function

The gills of vertebrates function in gas exchange, ion homeostasis, chemoreception and excretion of nitrogenous waste (Evans et al., 2005; Fu et al., 2010). The pseudobranch is often referred to as a non-respiratory structure in adult fishes, and given its irrigation by oxygenated blood and its relatively small size, it does seem unlikely that the pseudobranch contributes significantly to blood gas exchange. However, the pseudobranch may function in gas exchange during the early life history stages of some teleost fishes (Dunel, 1975, as cited in Laurent and Dunel-Erb, 1984). Given the remodelling of pseudobranch vasculature that occurs through ontogeny (see above), it seems plausible that the non-respiratory role of this structure reflects an evolutionary reduction following relaxed constraint and changing function of mandibular arch derivatives, rather than an independent origin of the pseudobranch to fulfil a physiological role distinct from that of the gills.

Indeed, our gene expression analyses lend support to hypotheses of generally shared function of gills and the pseudobranch. In tetrapods, GATA transcription factors function upstream of *Gcm2* to specify cells of the parathyroid gland, an endodermally-derived structure that functions in regulating extracellular calcium homeostasis (Kim et al., 1998; Günther et al., 2000; Gordon et al., 2001; Liu et al., 2007; Grigorieva et al., 2010). In fishes, this function is performed by the gills, which are also endodermally-derived (Warga and Nüsslein-Volhard, 1999; Gillis and Tidswell, 2017; Hockman et al., 2017), and which express *gcm2* and are generally regarded as homologues of the tetrapod parathyroid gland (Okabe and Graham, 2004). Our *in situ* gene expression data for *gata2, gata3 and gcm2* point to conservation of this function not only in the gills, but also in the mandibular arch pseudobranch. Additionally, the gills of fishes contain hypoxia-sensitive, chemoreceptive neuroepithelial cells (NECs), recognisable based on their morphology and co-expression of 5HT and SV2 (Dunel-Erb et al., 1982; Jonz & Nurse, 2003, 2005; Hockman et al., 2017). These cells are dispersed throughout gill lamellae, are sensitive to acute changes in partial pressure of oxygen, and elicit behavioural changes (e.g. increased breathing frequency and amplitude) to match oxygen supply to metabolic demand (reviewed by Milsom, 2012). Consistent with physiological evidence for a shared chemoreceptive function of the pseudobranch and gills (Laurent and Rouzeau, 1972), we find in skate that both contain 5HT/SV2+ putative hypoxia-sensitive NECs. Thus, contrary to arguments that the pseudobranch performs chemosensory or osmoregulatory functions that are distinct from those of the gills, our gene expression data are consistent with previous physiological studies in pointing to a broad conservation of pseudobranch and gill function in fishes.

### Serial homology of the spiracular cartilage and branchial rays of elasmobranch fishes

Each of the hyoid and gill arches of sharks, skates and rays are associated with a series of paired, cartilaginous appendages called branchial rays (Gillis et al., 2009b). These appendages articulate proximally with the epi- and ceratobranchial components of the arches, and project distally into interbranchial septa, where they serve as attachment points for branchiomeric musculature and support the respiratory lamellae of the gills. Branchial rays arise as discrete condensations that are distinct from their associated gill arch cartilages and develop under the influence of a *shh*-expression epithelial signalling centre called the gill arch epithelial ridge (GAER): shh signalling maintains proliferation of branchial ray progenitors, and progressively earlier inhibition of hedgehog signalling with the small molecule antagonist cyclopamine results in a successively greater reduction of the branchial ray series (Gillis and Hall, 2016). In cartilaginous fishes, the pseudobranch of the mandibular arch is supported by one or more spiracular cartilages that forms in close association with the palatoquadrate (upper jaw). In sharks, spiracular cartilages occur as multiple flattened rods that splay or bifurcate distally, while in skates the spiracular cartilage forms as a continuous sheet of cartilage (El-Toubi, 1947). It has been suggested that the spiracular cartilage could represent a mandibular arch homologue of the hyoid and gill arch branchial rays (Gegenbaur, 1872; El-Toubi, 1947; Holmgren and Stensio, 1936 as cited by El-Toubi, 1947) – or alternatively, a separate otic process of the palatoquadrate, which condenses as an extension of the latter, but then subsequently separates and chondrifies as a distinct element (Huxley, 1876; Parker, 1879; Edgeworth, 1925).

We find in skate that the spiracular cartilage condenses and chondrifies completely independently of the palatoquadrate, in a temporal and spatial configuration that parallels that of the branchial ray and gill arch cartilages. Moreover, we find that the spiracular cartilage develops beneath a *shh*-expressing GAER on the mandibular arch, from shh-responsive (i.e., *ptc*-expressing) mesenchyme, and that progressively earlier inhibition of hedgehog signalling with cyclopamine results in a successively greater reduction of the spiracular cartilage. These mandibular arch signalling interactions and spiracular cartilage phenotypes precisely mirror those of the hyoid and gill arch GAERs and branchial rays, and we therefore argue that the former are clear serial homologues of the latter (Fig. 5).

To date, branchial rays and spiracular cartilages have only been resolved to the chondrichthyan stem (Gillis et al., 2011; Burrow et al., 2020), and there is little fossil evidence of the presence of these features outside of chondrichthyans. Spiracles are often difficult to interpret in the fossil record. Some placoderms possessed an opening in the position of a spiracle (Young & Zhang, 1992; Arsenault et al., 2004), but distinguishing this feature from other gaps in their heavy dermal armour remains challenging. It is similarly challenging to reconstruct the internal anatomy of the spiracular region from fossil, though some have described grooves in stem gnathostome and stem osteichthyan skulls that may represent remnants of the efferent pseudobranchial artery (Goodrich, 1930; Gardiner, 1984; Basden & Young, 2001). A notable exception is the Middle Devonian acanthodiform *Cheiracanthus murchisoni* (Burrow et al., 2020). *Cheiracanthus* possessed a spiracular skeleton that takes the form of both rods and a flattened element in each spiracle, possibly forming a valve for controlled water flow over the pseudobranch. The series of thin parallel rods associated with the *Cheiracanthus* spiracular skeleton bears a strong resemblance to branchial rays and is more reminiscent of the rod-like spiracular cartilage of extant sharks than the sheet-like spiracular cartilage found in skate. *Cheiracanthus* may thus represent a transitional state between an ancestral gnathostome mandibular arch condition, with a less constrained first gill slit and a gill-arch-like mandibular arch, and the derived condition of extant elasmobranchs bearing spiracles with a reduced pseudobranch and supporting spiracular skeleton.

## Conclusion

Most questions of body plan evolution or anatomical novelty may be framed as hypotheses of historical and/or serial homology – i.e., homology of structures across or within taxa, respectively. Hypotheses of historical homology aim to resolve the polarity of character state changes on an evolutionary timescale by considering corresponding features within a phylogenetic context, while hypotheses of serial homology attempt to explain anatomical similarity due to shared underlying developmental mechanisms (Wagner, 1989, 1994; Panchen, 1992). But while hypotheses of serial homology may be formulated based on observation of anatomy or development even in a single taxon, such hypotheses must ultimately incorporate phylogeny, in order to distinguish between serial homology with ancestral polyisomery (repeated units that were ancestrally similar) or ancestral anisomery (repeated units that were ancestrally dissimilar, but that acquire a derived or secondary similarity) (Miyashita and Diogo, 2016).

The evolutionary history of the vertebrate mandibular arch remains obscure, owing to a dearth of fossils showing direct evidence of endoskeletal and/or soft tissue anatomy at the origin of jaws. We are therefore left in the somewhat difficult position of weighing limited, indirect evidence of ancestral mandibular arch anatomy and function from fossils against the striking anatomical and developmental similarities between the mandibular and gill arches of some extant taxa. In the case of the pseudobranch and spiracular cartilage, we can confidently infer that the former is an ancestral feature of the gnathostome mandibular arch, while the latter can (at present) only be resolved to the chondrichthyan stem.

Palaeontological consensus is that vertebrates likely possessed a non-respiratory mandibular arch, possibly resembling, anatomically, the oral apparatus of extant cyclostomes. We suggest, however, that the conserved, iterative development of the pseudobranch and gill lamellae of the mandibular, hyoid and gill arches (and, in chondrichthyans, their supporting skeletons) is too precisely similar in terms of embryology, gene expression and function to be attributable to convergent evolution. Furthermore, while there is currently no fossil evidence that that pseudobranch was ever a larger, more extensive mandibular arch gill, we suggest that arguments of co-option needn’t be invoked given the closely shared developmental context of these structures. Rather, we speculate that these similarities reflect an ancestral and deeply conserved gill arch-like nature of the gnathostome (if not vertebrate) mandibular arch, and that the pseudobranch of extant fishes is a vestige of this ancestral gill arch-like condition.

## Acknowledgements

We thank Dr. Richard Schneider, Prof. David Sherwood and the MBL Embryology course for provision of lab space, Louise Bertrand and Leica Microsystems for microscopy support and David Remsen, Scott Bennett, Dan Calzarette and staff of the Marine Biological Laboratory Marine Resources Center for technical and animal husbandry assistance. We also thank Dr. Kate Criswell, Dr. Vicky Sleight, Jenaid Rees, Dr. Kate Rawlinson, the University of Cambridge chordate evo-devo community and Dr. Carole Burrow for helpful discussion.

